# Large-scale, interpretable gene regulatory network inference through biologically informed matrix factorization

**DOI:** 10.64898/2026.07.15.738791

**Authors:** Soel Micheletti, Viola Fanfani, Julia Vogt, John Quackenbush, Jonas Fischer, Alexander Marx, Panagiotis Mandros

## Abstract

Gene regulatory networks (GRNs) provide a mechanistic framework for understanding how transcription factors coordinate gene expression to establish cellular identity and phenotype. Methods that integrate gene expression with motif-derived regulatory priors and other sources of biological information have substantially advanced gene regulatory network inference by reconstructing condition-specific regulatory architecture. These approaches estimate the evidence supporting regulatory interactions and have proven remarkably successful in a wide range of biological applications. A complementary view of regulatory networks, however, seeks to estimate the effect of those interactions on gene expression itself, providing a framework in which regulatory edges can be interpreted as activating or inhibitory influences on transcription.

We developed Giraffe, a biologically informed matrix factorization framework that jointly estimates transcription factor activities and gene regulatory networks by integrating gene expression, motif-based regulatory priors, and transcription factor protein-protein interactions. Giraffe estimates signed partial regulatory effects whose magnitude and sign can be interpreted as the strength and direction of transcriptional regulation. Building directly on the biological framework established by methods such as PANDA, Giraffe provides a complementary representation of gene regulatory networks that emphasizes mechanistic interpretation while remaining scalable, flexible, and computationally efficient.

Across synthetic benchmarks, six human tissues, yeast transcription factor perturbation experiments, and liver hepatocellular carcinoma, Giraffe accurately reconstructs regulatory interactions while distinguishing activating from inhibitory regulation with high accuracy. The inferred networks recover known features of tissue-specific regulation, correctly classify regulatory effects in transcription factor perturbation experiments, and identify biologically coherent changes in regulatory programs associated with liver cancer. Together, these results demonstrate that estimating the direction of transcriptional regulation provides a complementary perspective on gene regulatory networks that facilitates biological interpretation and hypothesis generation.

## 1 Introduction

Identifying gene regulatory mechanisms is important if we are to understand both how cellular processes are controlled in “normal” health and development, and altered as diseases develop, progress, and respond to therapies. Many methods use associations between genes based on their expression as a surrogate for regulation and search for “modules” of co-expressed genes; this is frequently done by computing the correlation [Langfelder and Horvath, 2008], partial correlation [Watson-Haigh et al., 2010], or mutual information [Faith et al., 2007, Meyer et al., 2007] based on gene expression levels across samples. Other methods infer Gene Regulatory Networks (GRNs), defined by relationships between transcription factors (TFs) and the genes they likely control. GRNs not only better represent the mechanics of the underlying biology, in which TFs bind to the promoter regions of their target genes to promote or inhibit their expression, but also have greater predictive power [Marbach et al., 2010, Lopes-Ramos et al., 2020].

Some GRN inference methods use regression, in which target gene expression is treated as the dependent variable and the expression levels of TFs that might regulate the gene are treated as the independent variables [Haury et al., 2012, Patel and Wang, 2015]. Other methods use machine learning frameworks such as message passing together with soft constraints based prior knowledge of likely TF-gene regulatory associations to infer condition-dependent GRNs starting from expression data [Glass et al., 2013, Weighill et al., 2022, Hossain et al., 2023]. Yet another set of methods extends these TF-gene association methods to also estimate TF activity defined as the degree to which TFs are available to regulate genes [Arrieta-Ortiz et al., 2015, Ma and Brent, 2021, Chen and Padi, 2024]. Unfortunately, the relationship between the expression and activity of a TF is complex, as many factors, including translation, protein modification, and degradation, can alter activity, and direct experimental measurement remains challenging.

Some models formulate estimating TF activity as a Bayesian matrix factorization [Chen and Padi, 2024, Gao et al., 2021], which makes it computationally intensive and requires assumptions for the posterior of the regulatory weights [Mar, 2019]. Such methods generally do not model interactions between TFs, which can miss true regulatory interactions. Others are based on network component analysis [Fu et al., 2011], multi-task learning [Castro et al., 2019], bilinear models [Ma and Brent, 2021], and variational inference [Mahmood et al., 2022]. However, these are often unable to scale beyond a few hundred genes and generally do not distinguish activating and inhibitory regulation, either because they incorporate complex nonlinear relationships that are challenging to interpret, or because they aim to infer scores reflecting confidence for the existence of regulatory relationships rather than magnitude and sign of association.

Many GRN inference methods estimate only the likelihood or confidence of regulatory interactions, without resolving whether a transcription factor activates or represses its targets. While such representations are useful for identifying putative regulatory relationships, they limit biological interpretation, as activation and repression have fundamentally different mechanistic and phenotypic consequences. Distinguishing these modes of regulation can shed light on pathway dynamics, help identify therapeutic targets, and aid interpretation of perturbation experiments. Despite its importance, directionality of regulation is rarely modeled explicitly in scalable genome-wide network estimation methods.

Giraffe, Gene-level Inference of Regulatory effects As Factorizations of Functions of Expressions, is a scalable method that integrates multiple sources of prior knowledge to jointly estimate an interpretable GRN and sample-specific transcription factor activities (Figure 1-A). Giraffe combines concepts from collaborative filtering [Koren et al., 2009, Wang et al., 2022] with biologically informed network inference in a matrix factorization framework. Collaborative filtering recommender systems are often framed as matrix factorization and have the advantage that they are designed to handle incomplete data [Kondratyeva et al., 2022]. Prior-informed approaches such as PANDA [Glass et al., 2013] and OTTER [Weighill et al., 2021] integrate information on likely TF-gene regulatory associations based on transcription factor binding motifs, protein-protein interactions between transcription factors, and gene expression to infer condition-specific regulatory networks, and have been demonstrated to perform well in a wide range of biological settings [Lopes-Ramos et al., 2018, Lopes-Ramos et al., 2020, Saha et al., 2023, Fanfani et al., 2024]. Giraffe builds on these biological principles to learn interpretable GRNs in which edge weights correspond to partial regulatory effects and, most importantly, edge signs distinguish activating from inhibitory regulation. Building on prior-informed approaches, Giraffe provides a complementary representation of gene regulatory networks that facilitates mechanistic interpretation while remaining scalable, flexible, and computationally efficient. We demonstrate Giraffe’s effectiveness using both synthetic and real-world data and then use it to explore hepatocellular carcinoma using data from TCGA.

**Figure 1:**
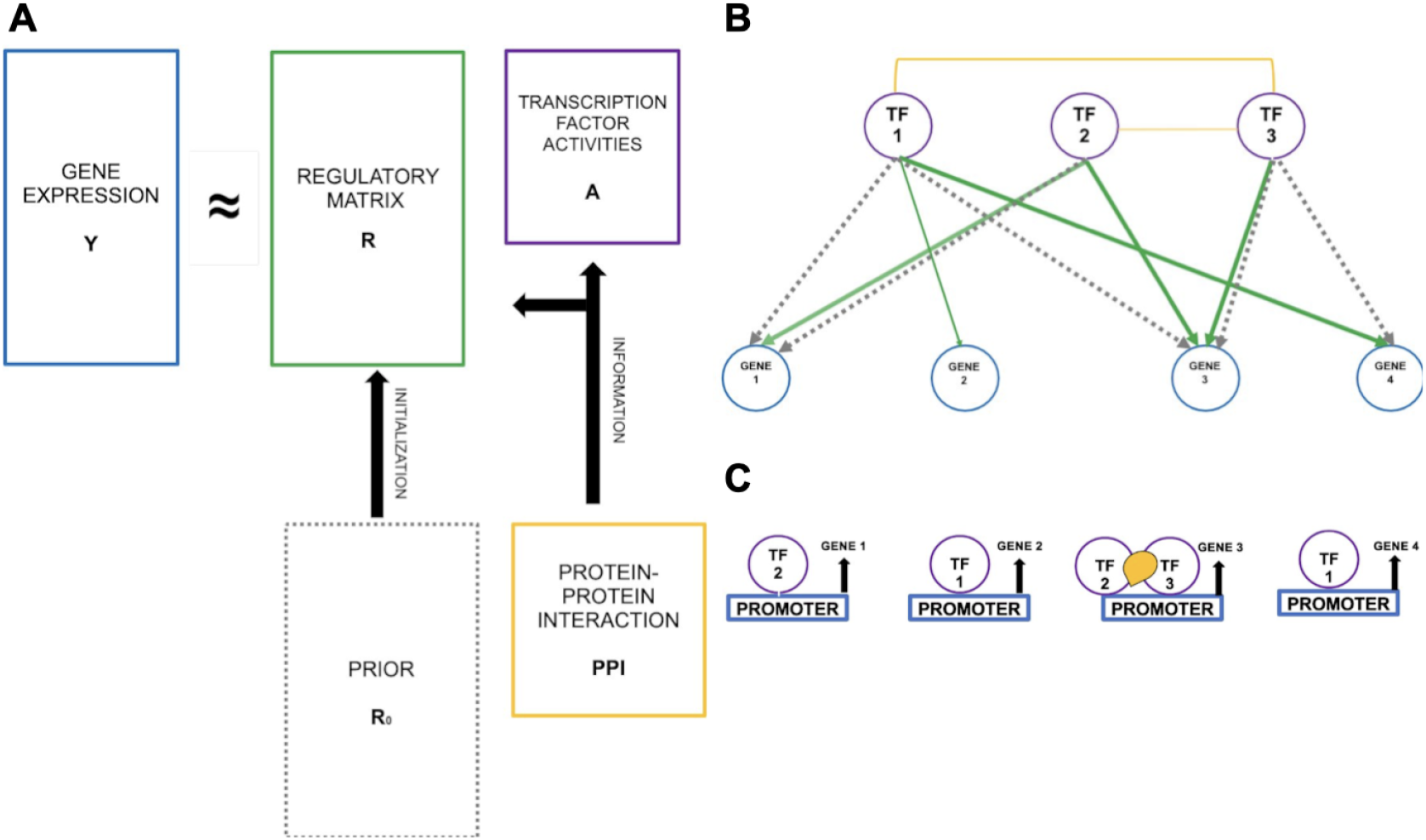
Joint estimation of regulation and transcription factor activities using Giraffe. (A) Schematic overview of Giraffe. Given gene expression by conditions matrix, *Y* , a prior matrix of likely TF-gene regulatory interactions, *R*_0_, and protein-protein interactions between TFs (PPI) data, Giraffe estimates a regulatory matrix *R* and a transcription factor activity matrix *A* through a biologically informed matrix factorization of *Y*. **(B) Our model of gene regulation**. A toy example with three TFs and four genes. Our goal is to estimate regulation, which is represented by directed edges (shown in bold green) from transcription factors to genes. First, protein-protein interactions are undirected edges between interacting transcription factors (orange solid edges) and capture the potential for regulatory complexes whose components may coordinately regulate a gene. Here, TF 1 and TF 3 have evidence of interaction, as do TF 2 and TF 3. Second, we also integrate a motif-based binary prior (gray dashed and green edges), which we use as an initial model of potential regulatory interactions; the green edges are those from the prior that the model estimates as “true” edges with evidence of regulatory activity. **(C) Gene regulation**. The model estimates the regulatory effects of transcription factors, including those acting as components of higher-order protein complexes, on the expression of their target genes.

## 2 Gene regulatory network inference with Giraffe

Giraffe decomposes a gene expression matrix *Y* as the product of a gene-to-transcription-factor matrix *R* and a non-negative transcription factor activity matrix *A* (Figure 1-B). Complementing methods that estimate the confidence that regulatory interactions exist, our formulation estimates the contribution of each transcription factor to target gene expression, allowing inferred regulatory effects to be interpreted directly as activating or inhibitory influences. By incorporating additional biological information, Giraffe yields an interpretable GRN that provides insight into TFs that are active regulators under various conditions.

### 2.1 Model

We model the regulation of genes by TFs as a graph *G* = (*V, E*) consisting of a set of nodes *V* = *V*_*G*_ ⊎ *V*_*T F*_ , representing genes and TFs, respectively, and a set of edges *E* ⊆ *V*_*G*_ × *V*_*T F*_ (Figure 1-B) that capture regulatory interactions. Our model incorporates two types of edges: directed TF-to-gene edges representing regulatory interactions and undirected TF–TF edges representing protein-protein interactions between transcription factors. In Giraffe, the goal is to infer the TF–gene regulatory edges. To guide this inference, we incorporate a motif-based prior for TF–gene pairs that captures the fact that not every TF has a binding site in the regulatory region of every gene (typically within a window surrounding the transcription start site). We also include a prior over TF–TF interactions, derived from protein-protein interaction data, to further constrain the inferred regulatory network.

Throughout the remainder of the manuscript, we let |*V*_*G*_|, |*V*_*T F*_ |, and *n* denote the number of genes, transcription factors, and biological samples, respectively. Gene expression is represented by the matrix 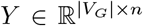, where rows correspond to genes and columns correspond to samples. We seek to approximate *Y* as the product of a regulatory effects matrix 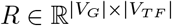 and a transcription factor activity matrix 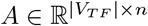. The entries of *R* represent signed partial regulatory effects of transcription factors on gene expression, while each column of *A* contains the latent transcription factor activities for an individual sample.

We model the regulatory relationship between transcription factors and genes using the linear model

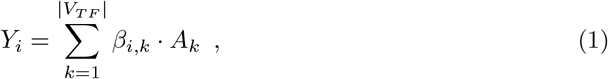

where *Y*_*i*_ ∈ ℝ^*n*^ is the expression profile of gene *i* across the *n* samples and *A*_*k*_ ∈ ℝ^*n*^ is the inferred activity profile of transcription factor *k* across those same samples. We chose this formulation because the coefficient *β*_*i,k*_ in Equation 1 can be interpreted as the partial effect of transcription factor *k* on the expression of gene *i*, after accounting for the activities of all other transcription factors in the model. Consequently, the inferred edge weights represent regulatory effects rather than simply evidence supporting regulatory interactions. The magnitude of each coefficient reflects the estimated strength of regulation, while its sign distinguishes activating (positive) from inhibitory (negative) effects.

Giraffe infers a GRN by estimating the coefficients *β*_*i,k*_ for all *i* ∈ [|*V*_*G*_|] and *k* ∈ [|*V*_*T F*_ |]. If the transcription factor activity matrix *A* were known, this would reduce to solving |*V*_*G*_| independent linear regression problems. In practice, however, transcription factor activities are generally not directly observable. While some studies have used transcription factor mRNA expression as a surrogate for activity [Ernst et al., 2008,Pierson et al., 2015], this approximation neglects translation, post-translational modification, protein degradation, and other processes that influence transcription factor activity, leading to potentially substantial discordance between TF expression and functional activity [Ma and Brent, 2021, Latchman, 1993]. Consequently, Giraffe jointly estimates the regulatory effects matrix *R* and the transcription factor activity matrix *A*, allowing the model to simultaneously infer regulatory relationships and latent transcription factor activities from the observed gene expression data. As we demonstrate throughout this work, this biologically informed formulation captures meaningful regulatory behavior while remaining computationally tractable and interpretable.

### 2.2 Objective function and optimization

We formulated Giraffe as a matrix factorization problem (Figure 1-A), where the gene expression matrix 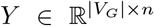 is decomposed as the product of a regulation matrix 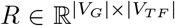 and a non-negative transcription factor activity matrix 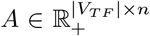. In most applications involving genome-wide gene expression data, the number of transcription factors (∼1600) is much larger than the number of samples, resulting in an underdetermined problem with an infinite number of solutions [Siegenthaler and Gunawan, 2014]. This can be seen in the matrix factorization formulation by setting *A* as a zero-padded identity matrix and *R* as a zero-padded copy of the gene expression matrix. This exactly reconstructs the gene expression, but does not reflect the underlying biology. Consequently, solely minimizing the reconstruction error is insufficient. To solve this optimization problem, we further constrain it by using two priors as soft constraints, each of which reflects distinct aspects of the regulatory process: a binary undirected graph 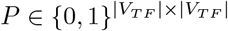 representing known protein-protein interactions (PPIs) between transcription factors, and a motif-based prior for the regulatory network 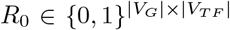 that maps regulatory TFs to their target genes such that an entry is one if and only if the sequence motif of the corresponding TF is found in the binding site of the target gene. The joint inference of *R* and *A* is achieved by solving the following optimization problem:

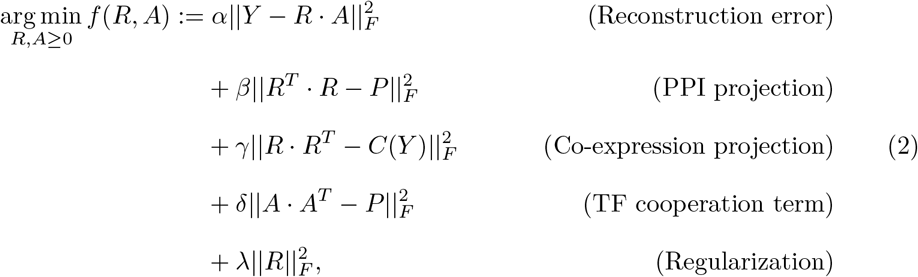

where || · ||_*F*_ is the Frobenius norm, and *C*(*Y*) is the co-expression matrix. The objective function in Equation 2 is a linear combination of multiple components. The PPI projection term encourages interacting proteins to target the same set of genes. Similarly, the TF cooperation term yields more correlated activities for interacting transcription factors. The co-expression projection term is based on the “guilt-by-association” principle [Chu et al., 1998] such that genes that are co-regulated are likely to be co-expressed. The final term shrinks the regression coefficients in a manner analogous to ridge regularization, mitigating the sensitivity to noise in the data and preventing overfitting.

To tune the hyperparameters *α, β, γ*, and *δ*, we balance the influence of the various inputs to enhance interpretability while minimizing the overall error. We set

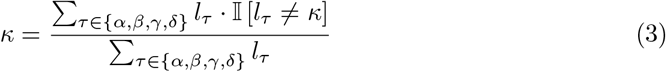

for *κ* ∈ {*α, β, γ, δ*}, where *l*_*τ*_ corresponds to the magnitude of the norm with coefficient *τ* ∈ {*α, β, γ, δ*} in *f*. Lee and Kim used a similar approach for weight initialization in the context of estimating depth in a single image [Lee and Kim, 2020]; this work demonstrated that the method effectively balances the contribution of each component in the loss function. We set *λ*=1 by default for all analyses presented here with the option of dynamically customizing the hyperparameters.

Minimizing *f* (*R, A*) is a non-convex problem, and we used the PyTorch implementation of the Adam optimizer [Kingma and Ba, 2014] with a learning rate of 10^−5^ and loss convergence as the stopping criterion. We initialized *R* using our motif-based prior as 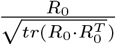 and *A* from a uniform distribution with support (0, 1).

Optimizing Giraffe’s objective as formulated in Equation 2 leads to a dense regulation matrix *R*, where all transcription factors are included in the set of parents of each target gene. To improve interpretability, our implementation includes the option of using the *ℓ*_1_-norm instead of the Frobenius norm in the regularization term of Equation 2. As in Lasso regression, this yields a sparse regulation matrix [Tibshirani, 1996]. Two considerations for this task are the use of the proximal operator in the optimizer (see [Melchior et al., 2019] and Appendix A in [Micheletti, 2023]), and the tuning of the regularization hyperparameter to address the underlying trade-off between type I and type II error. In our tutorial^1^, we show how *λ* can be picked via the stability selection framework [Meinshausen and Bühlmann, 2010]. Finally, since gene expression profiles may exhibit batch effects, we included two optional preprocessing steps in Giraffe: Limma [Ritchie et al., 2015] for location-scale correction and Cobra [Micheletti et al., 2024] for higher-order batch effect correction.

## 3 Results

We evaluated Giraffe using a progression of increasingly realistic settings. We began with *in silico* synthetic data, where the underlying regulatory network is known and network recovery can be quantified directly (Section 3.1). We then evaluated performance using gene expression data from six human tissues with independent ChIP-seq validation (Section 3.2), examined Giraffe’s ability to distinguish activating from inhibitory regulation using yeast transcription factor perturbation experiments (Section 3.3), and finally explored the biological insights provided by Giraffe in liver hepatocellular carcinoma (Section 3.4). Because the biological utility of a regulatory network ultimately depends on its ability to generate meaningful mechanistic hypotheses, we view these analyses as complementary, with increasing emphasis on experimentally supported and biologically interpretable results.

### 3.1 Improved recovery of regulatory interactions in *in silico* analyses

Synthetic benchmarks provide an important first assessment because the underlying regulatory network is known. At the same time, they necessarily depend on assumptions regarding network topology, transcription factor activity, and the mechanisms by which transcription factors regulate gene expression. Consequently, performance on simulated data should be viewed primarily as evidence that a method behaves as expected under a defined generative model rather than as definitive evidence of biological validity.

As described in Methods M1, we generated synthetic expression data resembling high-quality human transcriptomic data and inferred regulatory networks using Giraffe with motif priors exhibiting varying degrees of misspecification (“prior reliability”). We compared Giraffe with WGCNA [Langfelder and Horvath, 2008], Pearson correlation between TF and target gene expression (Cor), Tigress [Haury et al., 2012], Genie3 [Huynh-Thu et al., 2010], Bitfam [Gao et al., 2021], Tiger [Chen and Padi, 2024], Otter [Weighill et al., 2021], and Panda [Glass et al., 2013]. For each method we compared the inferred network with the known underlying regulatory network using AUROC. Results represent the mean and standard deviation over 50 independently generated datasets (Table 1, Tables S1–S3).

**Table 1:** Comparison of AUROC scores (rounded to two digits) for recovery of the underlying regulatory network topology by Giraffe and other GRN inference methods using synthetic data generated under varying levels of prior reliability. We report the mean and standard deviation across 50 independently generated datasets. Pearson correlation (Cor), WGCNA, Genie3, and Tigress have constant scores across prior reliability settings because they do not use prior biological information during network inference. These analyses evaluate recovery of network structure under the assumed generative model but do not assess the ability of a method to distinguish activating from inhibitory regulation. **Bold text** denotes the highest score for a given setting.

| AUROC | Prior reliability |  |  |  |  |
| --- | --- | --- | --- | --- | --- |
|  | 90% | 80% | 70% | 60% | 50% |
| GIRAFFE | **0.97** ± **0.001** | **0.93** ± **0.001** | **0.86** ± **0.002** | 0.78 ± 0.003 | 0.59 ± 0.004 |
| OTTER | 0.96 ± 0.004 | 0.91 ± 0.006 | 0.85 ± 0.008 | 0.74 ± 0.012 | 0.47 ± 0.004 |
| PANDA | 0.85 ± 0.022 | 0.83 ± 0.024 | 0.85 ± 0.021 | **0.84** ± **0.023** | 0.51 ± 0.030 |
| BITFAM | 0.53 ± 0.020 | 0.53 ± 0.020 | 0.53 ± 0.020 | 0.53 ± 0.020 | 0.53 ± 0.020 |
| TIGER | 0.59 ± 0.072 | 0.56 ± 0.065 | 0.56 ± 0.020 | 0.53 ± 0.019 | 0.50 ± 0.004 |
| GENIE3 | 0.66 ± 0.004 | 0.66 ± 0.004 | 0.66 ± 0.004 | 0.66 ± 0.004 | **0.66** ± **0.004** |
| TIGRESS | 0.58 ± 0.014 | 0.58 ± 0.014 | 0.58 ± 0.014 | 0.58 ± 0.014 | 0.58 ± 0.014 |
| COR | 0.58 ± 0.012 | 0.58 ± 0.012 | 0.58 ± 0.012 | 0.58 ± 0.012 | 0.58 ± 0.012 |
| WGCNA | 0.57 ± 0.006 | 0.57 ± 0.006 | 0.57 ± 0.006 | 0.57 ± 0.006 | 0.57 ± 0.006 |

Across simulations with prior reliability of at least 70%, Giraffe achieved the highest AUROC, while Panda and Otter consistently ranked among the strongest-performing methods. This result is noteworthy because all three methods leverage prior biological information, including transcription factor binding motifs and protein-protein interactions, suggesting that incorporating biological knowledge substantially improves network inference beyond expression data alone. In contrast, matrix factorization methods that do not explicitly incorporate TF-TF interactions (Bitfam and Tiger) performed considerably worse, particularly as the quality of the regulatory prior decreased. These results support the importance of modeling transcription factor cooperation during network inference.

The simulated datasets also recapitulate an important characteristic of biological systems: interacting transcription factors produce highly correlated regulatory activities, resulting in multicollinearity in gene expression [Salleh et al., 2017, Hikichi et al., 2020]. As expected, correlation-based approaches (Cor and WGCNA) performed only marginally better than random because they cannot effectively distinguish direct from indirect regulatory relationships under these conditions [Crow and Gillis, 2018]. Although regularization partially mitigates these effects, Tigress achieved performance similar to simple correlation analysis. By comparison, Genie3, which models nonlinear relationships without relying on prior biological information, performed best among methods that do not use TF-TF interaction data and was the strongest method when the regulatory prior became effectively uninformative (50% prior reliability). This behavior is expected because, at this level of misspecification, half of the entries in the motif prior have been randomized, removing most biologically meaningful information. Given the current state of transcription factor binding annotation, motif-derived priors are expected to be substantially more accurate than this worst-case scenario, making the higher prior reliability settings more representative of practical applications.

Importantly, these analyses evaluate only the recovery of network topology. They do not assess a principal feature of Giraffe: its ability to estimate the sign and magnitude of regulatory effects, thereby distinguishing activating from inhibitory regulation. That capability is evaluated using experimental perturbation data in the following sections, where biological interpretation becomes the primary focus.

### 3.2 Accurate GRN inference across human tissues

We next evaluated Giraffe using RNA-seq data from healthy individuals across six human tissues—breast, colon, kidney, liver, lung, and prostate—obtained from GRAND [Ben Guebila et al., 2022], comprising expression measurements for 30,243 protein-coding genes. We downloaded transcription factor protein-protein interactions from the STRING database [Szklarczyk et al., 2016]^2^, and motif-based TF-gene interactions from CIS-BP [Weirauch et al., 2014]; neither resource is tissue specific. For validation, we used tissue-specific ChIP-seq data from the hTFtarget database [Zhang et al., 2020] for breast, colon, liver, lung, and prostate, and from ReMap [Chèneby et al., 2018] for kidney.

Using these data, we inferred GRNs with Giraffe, Otter, Panda, Genie3, Tigress, and Bitfam. We also evaluated the unrefined binary motif prior as a biological baseline. Network accuracy was assessed by comparing inferred TF-gene interactions with tissue-specific ChIP-seq data using AUROC (Table 2, Figure S1).

**Table 2:** Comparison of runtime (hh:mm:ss) and AUROC for recovery of tissue-specific regulatory interactions inferred by Giraffe, Otter, Panda, Genie3, Tigress, and Bitfam across six human tissues using tissue-specific ChIP-seq data for validation. The binary motif prior is included as a biological baseline. Runtime reflects genome-scale network inference using 30,243 protein-coding genes. **Bold text** denotes the fastest runtime and highest AU-ROC for each tissue.

|  | Metrics | GIRAFFE | OTTER | PANDA | GENIE3 | TIGRESS | BITFAM | Motif Prior |
| --- | --- | --- | --- | --- | --- | --- | --- | --- |
| Breast | Runtime (hh:mm:ss) | **2:21** | 12:22 | 24:08 | 37:57 | 36:15 | 2:18:05 | - |
|  | Performance (AUROC) | **0.697** | 0.668 | 0.506 | 0.503 | 0.503 | 0.513 | 0.513 |
| Colon | Runtime (hh:mm:ss) | **2:03** | 22:15 | 20:17 | 35:03 | 30:10 | 2:18:03 | - |
|  | Performance (AUROC) | **0.651** | 0.562 | 0.508 | 0.501 | 0.505 | 0.524 | 0.515 |
| Kidney | Runtime (hh:mm:ss) | **3:58** | 16:56 | 18:35 | 12:10 | 19:21 | 46:17 | - |
|  | Performance (AUROC) | **0.565** | 0.547 | 0.535 | 0.500 | 0.502 | 0.521 | 0.533 |
| Liver | Runtime (hh:mm:ss) | **2:13** | 13:07 | 15:01 | 20:05 | 25:16 | 2:10:10 | - |
|  | Performance (AUROC) | **0.715** | 0.673 | 0.506 | 0.498 | 0.502 | 0.517 | 0.529 |
| Lung | Runtime (hh:mm:ss) | **2:22** | 11:47 | 24:13 | 67:15 | 25:56 | 2:05:20 | - |
|  | Performance (AUROC) | **0.730** | 0.474 | 0.535 | 0.501 | 0.501 | 0.518 | 0.505 |
| Prostate | Runtime (hh:mm:ss) | **2:22** | 14:45 | 15:06 | 15:48 | 28:31 | 1:41:32 | - |
|  | Performance (AUROC) | **0.780** | 0.511 | 0.545 | 0.500 | 0.503 | 0.517 | 0.562 |

The results, summarized in Table 2, further emphasize the importance of incorporating biological prior knowledge during network inference. The motif prior alone outperformed Genie3, Tigress, and Bitfam across all tissues, indicating that experimentally and computationally derived information on transcription factor binding provides a strong foundation for constructing tissue-specific regulatory networks. Methods that integrate these priors with gene expression data, including Panda, Otter, and Giraffe, consistently achieved the highest overall performance. These findings reinforce the importance of combining transcriptomic measurements with biological knowledge rather than relying on gene expression alone.

Among the prior-informed approaches, Giraffe consistently achieved the highest AU-ROC across all six tissues, while Panda and Otter remained close competitors. Matrix factorization alone was not sufficient to achieve comparable performance, as Bitfam, despite also jointly estimating transcription factor activities and regulatory networks, performed only marginally better than the motif prior. Together, these observations suggest that modeling transcription factor cooperation through protein-protein interactions substantially improves the reconstruction of tissue-specific regulatory architecture.

Although the AUROC values are lower than would be expected for many conventional machine learning tasks, they are consistent with previous studies demonstrating that genome-scale GRN inference from noisy human transcriptomic data remains an exceptionally challenging problem [Zhao et al., 2021, Chen and Mar, 2018, Weighill et al., 2021]. In this context, the consistent improvement achieved by Giraffe across multiple tissues is notable, particularly in the lung and prostate datasets where several competing methods performed close to random. Giraffe also showed the largest improvement over the motif prior among the prior-informed methods, suggesting that it effectively combines motif information, transcription factor cooperation, and gene expression to better capture tissue-specific patterns of regulation. In addition, Giraffe completed inference at least fivefold faster than the competing methods, making it practical for genome-scale analyses involving tens of thousands of genes and large patient cohorts.

While these analyses demonstrate improved recovery of tissue-specific regulatory interactions, they evaluate only whether regulatory interactions are correctly identified. They do not assess one of the principal features of Giraffe: that the inferred edge weights correspond to signed partial regulatory effects whose magnitude and sign can be interpreted as the strength and direction of transcriptional regulation. We therefore next examined this capability directly using experimental transcription factor perturbation data.

### 3.3 Distinguishing activating from inhibitory regulation in yeast

A central feature of Giraffe is that its edge weights represent estimated partial regulatory effects, allowing both the magnitude and sign of each interaction to be interpreted biologically. Complementing methods designed to estimate the confidence that a regulatory interaction exists, Giraffe estimates whether a transcription factor is predicted to activate or repress expression of its target gene. To evaluate this capability, we analyzed *Saccharomyces cerevisiae* expression data for 778 genes following knockout of 42 individual transcription factors (TFKO) [Ma and Brent, 2021] (Methods M2).

If the expression of gene *i* increased following knockout of transcription factor *j*, relative to its average expression across the remaining experiments, we interpreted this as evidence that TF *j* normally represses gene *i*. Conversely, reduced expression following knockout was interpreted as evidence that TF *j* normally activates gene *i*. These perturbation experiments therefore provide an experimentally grounded estimate of regulatory direction, allowing us to assign each TF-gene interaction as activating or inhibitory. Approximately 48% of interactions were classified as activating and 52% as inhibitory.

We applied Giraffe using the gene expression data together with a ChIP-seq-derived binary regulatory prior [Ma and Brent, 2021] and transcription factor protein-protein interactions from STRING [Szklarczyk et al., 2016]. The inferred networks achieved a mean sign accuracy of 83%, ranging from 68% to 90% across the 42 transcription factor perturbations (Figure 2). We compared these results with linear regression, ridge regression (using TF expression to predict target gene expression), Pearson correlation (Cor), Tigress, Bitfam, and Tiger. Across virtually all perturbations, Giraffe predicted the direction of regulation substantially more accurately than the competing approaches, exceeding the next best-performing method by at least 20 percentage points (Figure 2).

**Figure 2:**
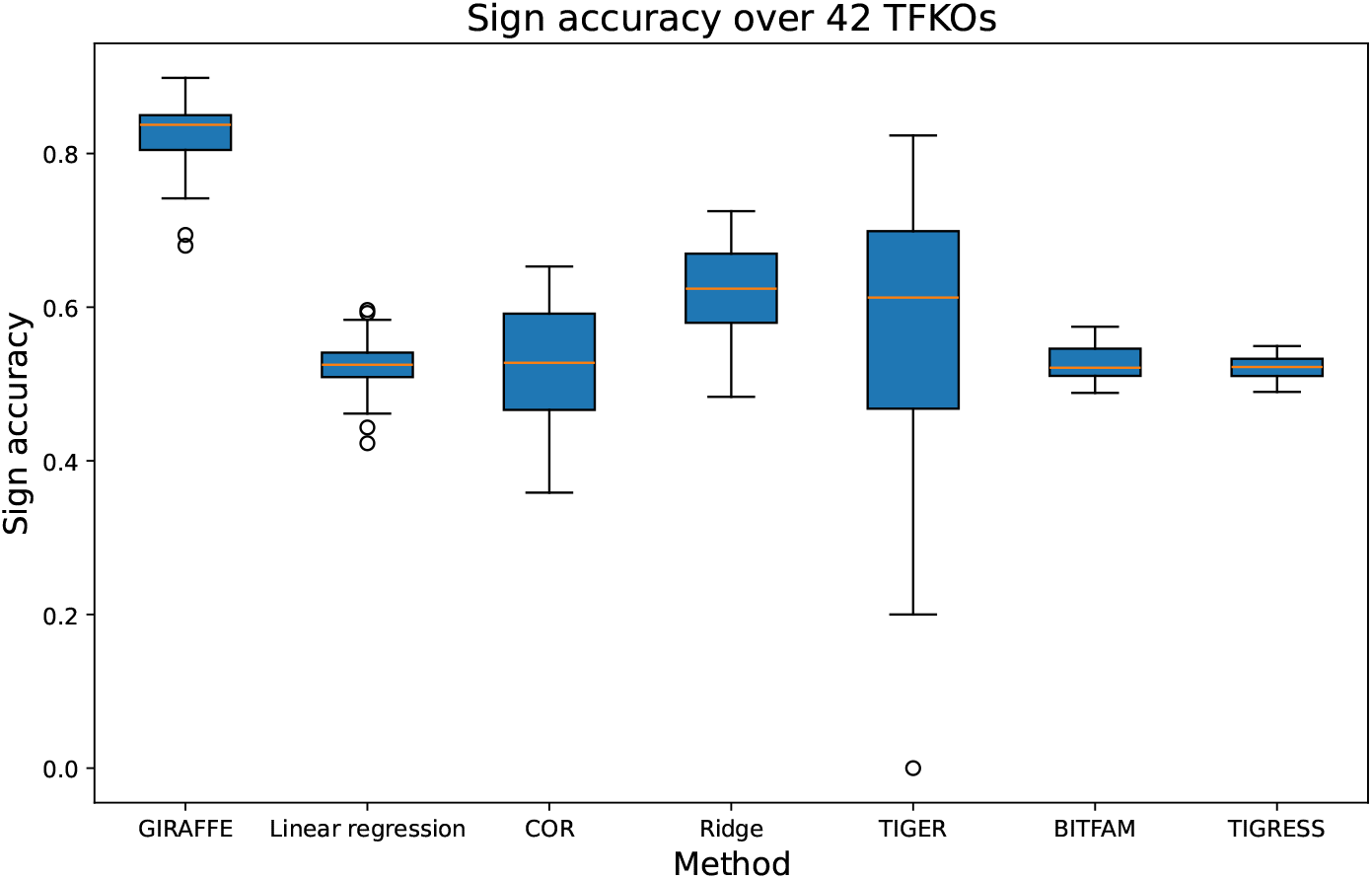
Experimental validation of signed regulatory effects using yeast transcription factor perturbations. Distribution of the accuracy with which each method distinguishes activating from inhibitory TF-gene regulatory interactions across 42 *Saccharomyces cerevisiae* transcription factor knockout experiments. For each method, bars show the median, interquartile range, whiskers, and extreme values of the sign accuracy across the 42 knockouts (778 genes). Giraffe consistently achieves the highest sign accuracy, demonstrating that its estimated partial regulatory effects capture biologically meaningful regulatory directionality.

These results demonstrate that Giraffe not only reconstructs regulatory network structure, but also estimates biologically meaningful signed regulatory effects. This complementary representation of gene regulatory networks provides mechanistic information that cannot be obtained from methods whose edge weights represent only the confidence that an interaction exists.

### 3.4 Analysis of regulation in liver hepatocellular carcinoma

We also used Giraffe to infer GRNs in liver hepatocellular carcinoma (LIHC) using TCGA data. Following the preprocessing of [Micheletti et al., 2024], the resulting dataset had gene expression values for 19674 genes and 424 samples–374 “cases” (samples classified as metastatic and primary tumor) and 50 “controls” (normal tissue adjacent to tumors, NAT). We used Giraffe to infer case and control GRNs with motif-based TF-gene priors [Weirauch et al., 2014] and protein-protein interaction [Szklarczyk et al., 2016] as prior information. Following GRN inference, we performed a graph differential analysis between cases and controls using node2vec2rank (nvr) [Mandros et al., 2024], which estimates a joint node embedding and ranks genes by their relative distances between conditions in the latent space. We used the nvr rank-ordered genes as input to Gene Set Enrichment Analysis (GSEA) implemented in the *prerank_gseapy* package (KS test, *p <* 0.05 with Benjamini-Hochberg FDR correction) and used the Kyoto Encyclopedia of Genes and Genome (KEGG) pathway database as the annotation source.

As expected, we found several biological pathways typically associated with liver cancer (Figure 3) to be differentially regulated based on the Giraffe GRN models. This includes pathways involving the cytoplasmic intermediates mTOR, MAPK signaling, PI3K, and their mediator phosphatidylinositol, all of which have been reported to be frequently altered in LIHC and are key targets for therapeutic intervention [Garcia-Lezana et al., 2021, Zhou et al., 2011]. The ErbB signaling pathway–which is crucial for the regulation of cell proliferation, migration, and apoptosis–influences LIHC progression and intersects with the PI3K/Akt, JAK/STAT, and MAPK pathways [Ho et al., 2017, Stefani et al., 2021]. Correct organization of actin cytoskeleton, which has been shown to be associated with adherens junction [Gerber et al., 2022], is vital in the liver homeostasis and disease control [Duwe et al., 2022]. Enriched pathways also include fatty acid metabolism, which has recently been reported as a potential target for liver cancer therapy [Wu and Lin, 2024], insulin signaling, which is also known to be disregulated in LIHC [Sakurai et al., 2017], and others such as those annotated as tight junction, progesterone-mediated oocyte maturation, endocytosis, and ABC transporters which have been shown to be involved in other cancers, including those of the liver [Lee and Luk, 2010, Chen et al., 2018, Blondy et al., 2019]. Other enriched pathways are associated with cancer and overall disease severity and include those annotated as pathways in cancer, GnRH (gonadotropin-releasing hormone), long-term potentiation, ubiquitin-proteasome system, ECM receptor, and focal adhesion [Gründker and Emons, 2017, Wang et al., 2023, Popova and Jücker, 2022, Provenzano and Keely, 2009]. As a comparison, a similar analysis using Bitfam identified fewer pathways and included generally less relevant terms such as general associations with various cancers and Leishmania infection (Figure S2).

**Figure 3:**
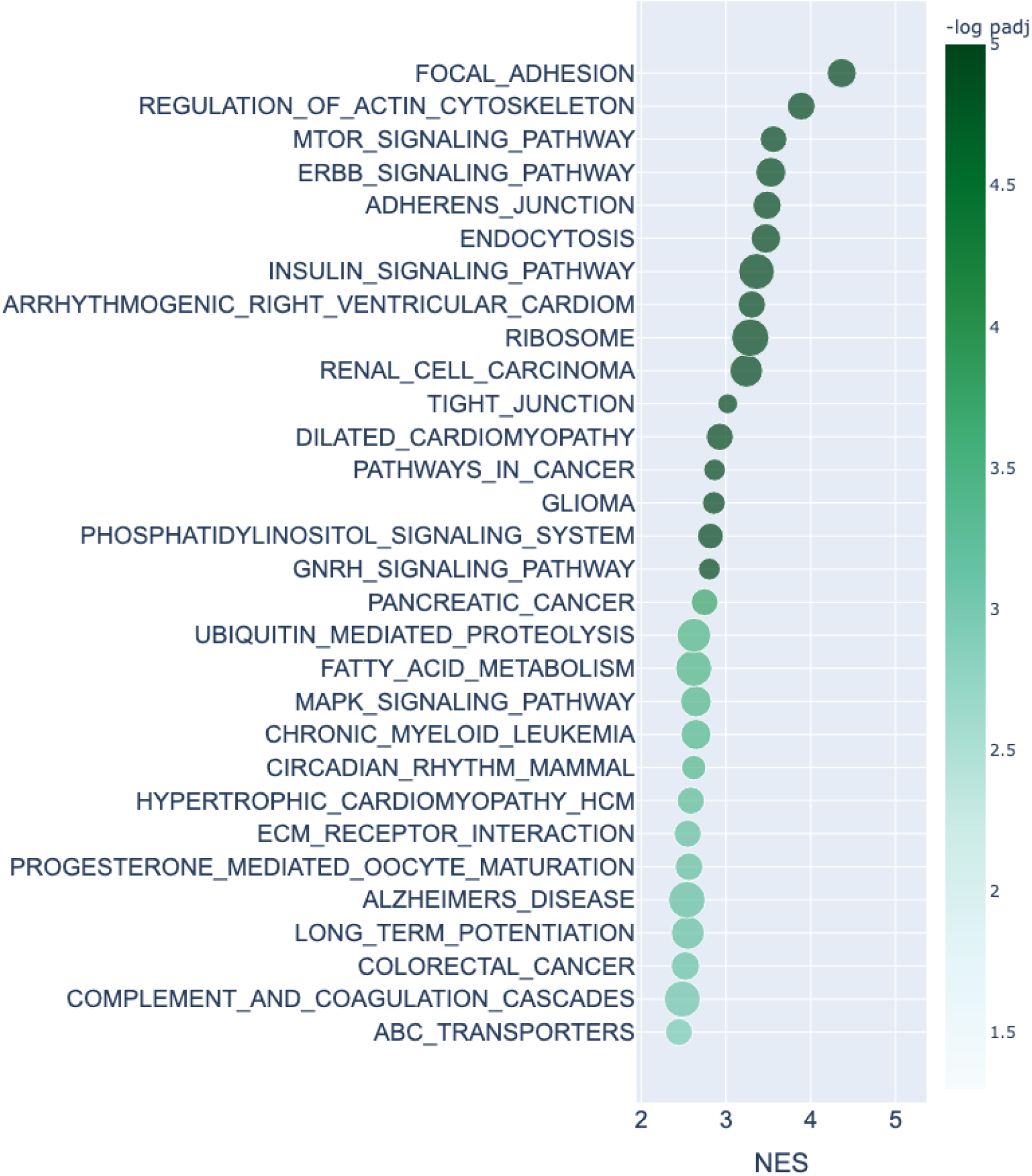
Differentially rewired biological pathways identified from Giraffe regulatory networks in liver hepatocellular carcinoma. KEGG pathway enrichment analysis of genes ranked by differential network topology between hepatocellular carcinoma and normal adjacent liver tissue.

Notably, the signed nature of the Giraffe networks enables us to distinguish between pathways that are differentially activated versus those that are differentially repressed in LIHC. For example, pathways such as PI3K/Akt and MAPK signaling are not only identified as dysregulated, but their inferred regulatory shifts suggest increased activation consistent with their known oncogenic roles. Conversely, pathways associated with normal liver function and metabolic homeostasis show patterns consistent with loss of regulatory activation or increased repression. This distinction provides a more nuanced view of tumor biology, as it separates pathway engagement from the direction of regulatory control, offering insights into potential mechanisms of disease progression and therapeutic intervention that are not accessible in unsigned network models.

## 4 Discussion

Although biological states are often characterized by patterns of gene expression, it is the regulatory processes that generate those expression patterns that ultimately define phenotype. Gene regulatory network (GRN) inference methods seek to reconstruct these regulatory relationships, typically representing each edge as the likelihood or confidence that a transcription factor regulates a target gene. Such representations have proven to be remarkably successful in identifying biologically meaningful regulatory networks, but generally do not distinguish whether a transcription factor activates or represses the expression of its targets. Because activation and repression have fundamentally different biological consequences, estimating the effects of regulatory interactions provides a complementary perspective for interpreting transcriptional programs.

Like other biologically informed approaches, including message-passing methods such as Panda, Giraffe uses prior biological knowledge to constrain an otherwise underdetermined inference problem. Motif-derived regulatory priors and transcription factor protein-protein interaction networks substantially reduce the solution space, guiding inference toward biologically plausible regulatory networks. Building on this framework, Giraffe jointly estimates transcription factor activities and regulatory effects, allowing network edges to be interpreted as signed partial regulatory effects on gene expression while retaining scalability to genome-wide datasets.

Because a definitive “ground truth” for gene regulatory networks is generally unavailable, we deliberately evaluated Giraffe using several complementary benchmarking strategies, including controlled *in silico* simulations, validation against tissue-specific ChIP-seq data, transcription factor perturbation experiments in yeast, and differential network analysis of liver hepatocellular carcinoma.

Although synthetic benchmarking provides a useful controlled environment for comparing methods, it necessarily depends on assumptions regarding regulatory architecture and therefore cannot fully capture the complexity of biological systems. Within this setting, Giraffe consistently recovered the underlying regulatory network more accurately than competing approaches whenever the regulatory prior reflected realistic levels of biological knowledge. More broadly, these experiments reinforced the importance of incorporating prior biological information into GRN inference. Methods that integrated motif-derived regulatory priors consistently outperformed those relying solely on gene expression, supporting the idea that domain knowledge remains an essential input for accurately reconstructing genome-scale regulatory networks.

The analyses using human tissue data provide a more biologically relevant assessment of performance. Across six healthy tissues, Giraffe consistently outperformed competing methods when evaluated against tissue-specific ChIP-seq data while remaining substantially faster than alternative prior-informed approaches. These results demonstrate that biologically informed GRN inference can achieve improved accuracy without sacrificing computational scalability, an an important consideration for the analysis of large transcriptomic datasets.

Analysis of yeast transcription factor knockout experiments further demonstrated that Giraffe accurately distinguishes activating from inhibitory regulatory interactions. This capability represents an important extension of genome-scale GRN inference because it provides information beyond network topology alone. Similarly, differential analysis of hepatocellular carcinoma identified regulatory rewiring in pathways central to liver cancer biology while providing a framework for interpreting changes in regulatory direction in addition to changes in network connectivity. Together, these analyses illustrate how signed regulatory effects provide an additional layer of biological interpretation beyond identifying whether a regulatory interaction exists.

Overall, Giraffe provides a complementary representation of gene regulatory networks that extends the growing family of biologically informed methods for GRN inference. Like Panda and related approaches, it integrates motif-derived regulatory priors and transcription factor protein-protein interactions to infer condition-specific regulatory networks while remaining scalable to genome-wide studies. Because Giraffe is formulated as a biologically informed matrix factorization framework, it also provides a natural foundation for future extensions, including GPU acceleration, single-cell applications, longitudinal analyses, and perturbation-based studies.

Rather than asking only whether a transcription factor regulates a target gene, Giraffe also estimates how that regulator influences gene expression once it acts, providing a complementary representation of gene regulatory networks in which regulatory edges can be interpreted as signed partial regulatory effects. We expect this complementary perspective to prove particularly valuable in studies of development, disease progression, therapeutic response, and perturbational biology, where changes in regulatory direction are often as important as changes in network topology. Rather than asking only whether a transcription factor regulates a target gene, Giraffe also estimates how that regulator influences gene expression once it acts, providing a complementary representation of regulatory networks that links network structure to regulatory mechanism.

## Code and data availability

Giraffe is available in Python through the netZoo package [Ben Guebila et al., 2023] netZooPy v0.11.0 (https://netzoo.github.io). We provide a tutorial, the code to reproduce our analyses, and a full version of the Master’s thesis that served as the foundation for this work [Micheletti, 2023] on GitHub (https://github.com/soelmicheletti/giraffe). Processed input data has been deposited in Zenodo (https://zenodo.org/record/7852640#.ZEKfV5FBxkg).

## Acknowledgements

The results shown here are in part based upon data generated by the TCGA Research Network: https://www.cancer.gov/tcga. The authors thank the Quackenbush group members for constructive feedback. SM gratefully acknowledges the Swiss Study Foundation for providing enriching opportunities and supporting his Master’s thesis. SM, JQ, JF, and PM were supported by grants from the US National Institutes of Health R35CA220523 and R01HG011393. During the project, AM was supported by a postdoctoral fellowship from the ETH AI Center.

## Methods

### M1. *In silico* synthetic data generation

To benchmark GRN inference under controlled conditions in which the underlying regulatory network is known, we generated synthetic expression data designed to capture key features of genome-scale regulatory systems while allowing direct comparison between inferred and ground-truth regulatory networks. We generated a synthetic dataset with *n* = 50 samples in which we modeled |*V*_*G*_| = 500 genes and |*V*_*T F*_ | = 100 transcription factors (additional configurations are available in the Supplementary Materials). We chose |*V*_*G*_| *>* |*V*_*T F*_ | *> n* to reflect the relative scale encountered in biological systems. For these networks, we generated the ground truth regulatory effects matrix 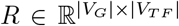 and transcription factor activity matrix 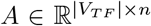, together with Giraffe’s input protein-protein interaction network 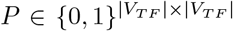, gene expression matrix 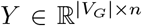, and motif prior 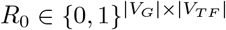.

Because Giraffe is based on the assumption that TF–TF interactions can result in protein complexes that regulate gene expression, we generated a symmetric network *P* using the Barabasi-Albert model (an algorithm for generating random scale-free networks using a preferential attachment mechanism) [Albert and Barabási, 2002], with interaction probabilities estimated from the STRING database on human data [Szklarczyk et al., 2016]. This ensured that the simulated PPI network exhibited the power-law degree distribution characteristic of biological protein-protein interaction networks [Wuchty et al., 2006]. We modeled transcription factor complexes by assuming that each densely connected PPI community (or “clique”) represents a regulatory complex capable of coordinately regulating common target genes. We then used these complexes to generate the ground-truth regulatory network *R* and the transcription factor activity matrix *A*.

For each *k* ∈ *K*, where *K* is the set of cliques in *P* , we generated a sample-specific activity vector *A*_*k*_ ∈ ℝ^*n*^ sampled independently from *U* (0, 1) together with a sparse regulatory vector 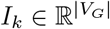 whose non-zero entries were sampled independently from 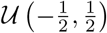. The sparsity of 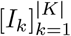 reflects the biological observation that transcription factors and transcription factor complexes regulate only a relatively small subset of genes. Based on these assumptions, we constructed the ground-truth transcription factor activity matrix *A* and regulatory effects matrix *R*, for *i* ∈ [|*V*_*T F*_ |], as

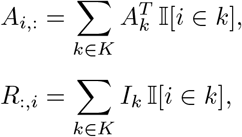

Finally, we generated the simulated gene expression matrix as *Y*:= *R* · *A*.

In its network inference model, Giraffe uses a motif prior, *R*_0_, that maps transcription factors to genes they have the potential to regulate based on the presence of transcription factor binding motifs within putative promoter regions. To assess the performance of Giraffe on simulated data, we generated a binary motif prior by setting all non-zero entries of *R* equal to one. Because motif-based priors are imperfect in biological systems—the presence of a transcription factor binding site does not necessarily imply binding or regulation—we introduced varying levels of prior misspecification by randomly flipping entries from 0 to 1 (or vice versa). Throughout the manuscript, we define the “prior reliability” as the fraction of entries that remain unchanged. Thus, a prior in which 10% of entries have been flipped is described as having a prior reliability of 90%.

### M2. Assessing network inference and method performance metrics

We compared the accuracy and performance of Giraffe with Tigress [Haury et al., 2012], Genie3 [Huynh-Thu et al., 2010], Bitfam [Gao et al., 2021], Otter [Weighill et al., 2021], and Panda [Glass et al., 2013]. For the synthetic data analyses, we additionally included WGCNA [Langfelder and Horvath, 2008], Pearson correlation between transcription factor and target gene expression (Cor), and Tiger [Chen and Padi, 2024]. We assessed computational efficiency by measuring inference runtime, with all methods executed on the same computational architecture [Lee and Tzou, 2009].

We assessed the quality of the inferred networks using the area under the receiver operating characteristic curve (AUROC). For simulated data, we compared the inferred networks with the known ground-truth regulatory network used to generate the expression data. Assessing network accuracy for human and yeast data is considerably more difficult because the true regulatory networks are unknown. We therefore used experimentally determined transcription factor binding from condition-specific ChIP-seq data, which measure TF–DNA interactions, as an independent validation resource wherever available. We evaluated only TF–gene pairs for which ChIP-seq evidence existed and treated GRN inference as a binary classification problem. Because this benchmark evaluates recovery of regulatory interactions rather than their direction, we compared the absolute values of the inferred partial regulatory effects. In this setting, both large positive and large negative coefficients provide strong evidence for the existence of a regulatory interaction.

Human genome-wide expression datasets typically include more than 25,000 genes, making inference computationally demanding for several methods. To enable comparison, we partitioned the data into subsets, inferred subnetworks independently, and aggregated them into complete GRNs. We used a batch size of 5000 genes for Tigress and 3000 genes for Bitfam. For Genie3, we reduced the default number of trees from 100 to 5 to obtain practical runtimes.

To evaluate Giraffe’s ability to distinguish activating from inhibitory regulation, we used the processed yeast perturbation dataset reported by [Chen and Padi, 2024], which includes ChIP-seq data together with genome-wide expression measured at baseline, following transcription factor knockout (TFKO), and following transcription factor overexpression (TFOE). We compared the signs inferred by Giraffe with a reference derived from the TFKO experiments, interpreting a decrease in expression of gene *j* following knockout of transcription factor *i* (relative to the unperturbed expression) as evidence that TF *i* positively regulates gene *j*, and an increase in expression as evidence that TF *i* negatively regulates gene *j*. Because these perturbation experiments directly alter transcription factor activity, they provide one of the few experimental settings in which the inferred direction of regulation can be evaluated.

## Footnotes

1 https://github.com/soelmicheletti/giraffe/blob/main/Tutorial.ipynb

2 https://github.com/soelmicheletti/giraffe/blob/main/notebooks/data/colon/preprocessing.py contains a sample script with detailed parameters.

## Notes

https://netzoo.github.io

https://github.com/soelmicheletti/giraffe

https://zenodo.org/record/7852640#.ZEKfV5FBxkg

### Competing Interest Statement

The authors have declared no competing interest.

