## Supplementary Materials for "Large-scale, interpretable gene regulatory network inference through biologically informed matrix factorization"

Micheletti et al.

### 1 Overview

This Supplement provides additional benchmarking analyses supporting the analyses presented in the main text. These include synthetic benchmark experiments across multiple sample sizes, complete ROC curves for tissue-specific ChIP-seq validation, and a comparison of pathway enrichment obtained using Bitfam in the liver hepatocellular carcinoma analysis.

### 2 Performance on synthetic datasets with varying sample sizes

To assess the robustness of Giraffe to sample size, we repeated the synthetic benchmarking analyses described in the main text using datasets containing 10, 25, and 75 samples. The overall trends remained consistent across all settings.

| **n=25** | Prior reliability | | | | |
|---|---|---|---|---|---|
| AUROC | 90% | 80% | 70% | 60% | 50% |
| GIRAFFE | **0.97 ± 0.001** | **0.93 ± 0.001** | **0.86 ± 0.001** | 0.76 ± 0.003 | 0.53 ± 0.006 |
| OTTER | 0.96 ± 0.004 | 0.91 ± 0.006 | 0.85 ± 0.008 | 0.74 ± 0.012 | 0.47 ± 0.004 |
| PANDA | 0.86 ± 0.009 | 0.86 ± 0.026 | 0.86 ± 0.016 | **0.85 ± 0.026** | 0.50 ± 0.028 |
| GENIE3 | 0.59 ± 0.002 | 0.59 ± 0.002 | 0.59 ± 0.002 | 0.59 ± 0.002 | **0.59 ± 0.002** |
| COR | 0.56 ± 0.010 | 0.56 ± 0.010 | 0.56 ± 0.010 | 0.56 ± 0.010 | 0.56 ± 0.010 |
| WGCNA | 0.54 ± 0.005 | 0.54 ± 0.005 | 0.54 ± 0.005 | 0.54 ± 0.005 | 0.54 ± 0.005 |

Table S1: Comparison of AUROC scores (rounded to two decimal places) for GRNs inferred by Giraffe and comparison methods using synthetic benchmark datasets with $n = 25$ samples. Results are reported as mean ± standard deviation across 50 independently generated datasets. Boldface indicates the best-performing method for each level of prior reliability.

| **n=75** AUROC | Prior reliability 90% | 80% | 70% | 60% | 50% |
|---|---|---|---|---|---|
| GIRAFFE | **0.97 ± 0.005** | **0.93 ± 0.001** | **0.87 ± 0.001** | 0.78 ± 0.002 | 0.58 ± 0.004 |
| OTTER | 0.96 ± 0.001 | 0.91 ± 0.006 | 0.85 ± 0.010 | 0.74 ± 0.012 | 0.47 ± 0.005 |
| PANDA | 0.86 ± 0.020 | 0.87 ± 0.019 | 0.85 ± 0.013 | **0.87 ± 0.012** | 0.50 ± 0.047 |
| GENIE3 | 0.72 ± 0.010 | 0.72 ± 0.010 | 0.72 ± 0.010 | 0.72 ± 0.010 | **0.72 ± 0.010** |
| COR | 0.59 ± 0.016 | 0.59 ± 0.016 | 0.59 ± 0.016 | 0.59 ± 0.016 | 0.59 ± 0.016 |
| WGCNA | 0.59 ± 0.007 | 0.59 ± 0.007 | 0.59 ± 0.007 | 0.59 ± 0.007 | 0.59 ± 0.007 |

Table S2: Comparison of AUROC scores (rounded to two decimal places) for GRNs inferred by GIRAFFE and comparison methods using synthetic benchmark datasets with $n = 75$ samples. Results are reported as mean ± standard deviation across 50 independently generated datasets. Boldface indicates the best-performing method for each level of prior reliability.

| **n=10** AUROC | Prior reliability 90% | 80% | 70% | 60% | 50% |
|---|---|---|---|---|---|
| GIRAFFE | 0.92 ± 0.005 | 0.85 ± 0.004 | 0.80 ± 0.006 | 0.70 ± 0.009 | 0.52 ± 0.008 |
| OTTER | **0.96 ± 0.004** | **0.91 ± 0.006** | 0.85 ± 0.008 | 0.74 ± 0.012 | 0.47 ± 0.004 |
| PANDA | 0.85 ± 0.022 | 0.83 ± 0.022 | **0.87 ± 0.024** | **0.85 ± 0.031** | 0.49 ± 0.034 |
| GENIE3 | 0.53 ± 0.002 | 0.53 ± 0.002 | 0.53 ± 0.002 | 0.53 ± 0.002 | **0.53 ± 0.002** |
| COR | 0.53 ± 0.006 | 0.53 ± 0.006 | 0.53 ± 0.006 | 0.53 ± 0.006 | 0.53 ± 0.006 |
| WGCNA | 0.51 ± 0.005 | 0.51 ± 0.005 | 0.51 ± 0.005 | 0.51 ± 0.005 | 0.51 ± 0.005 |

Table S3: Comparison of AUROC scores (rounded to two decimal places) for GRNs inferred by GIRAFFE and comparison methods using synthetic benchmark datasets with $n = 10$ samples. Results are reported as mean ± standard deviation across 50 independently generated datasets. Boldface indicates the best-performing method for each level of prior reliability.

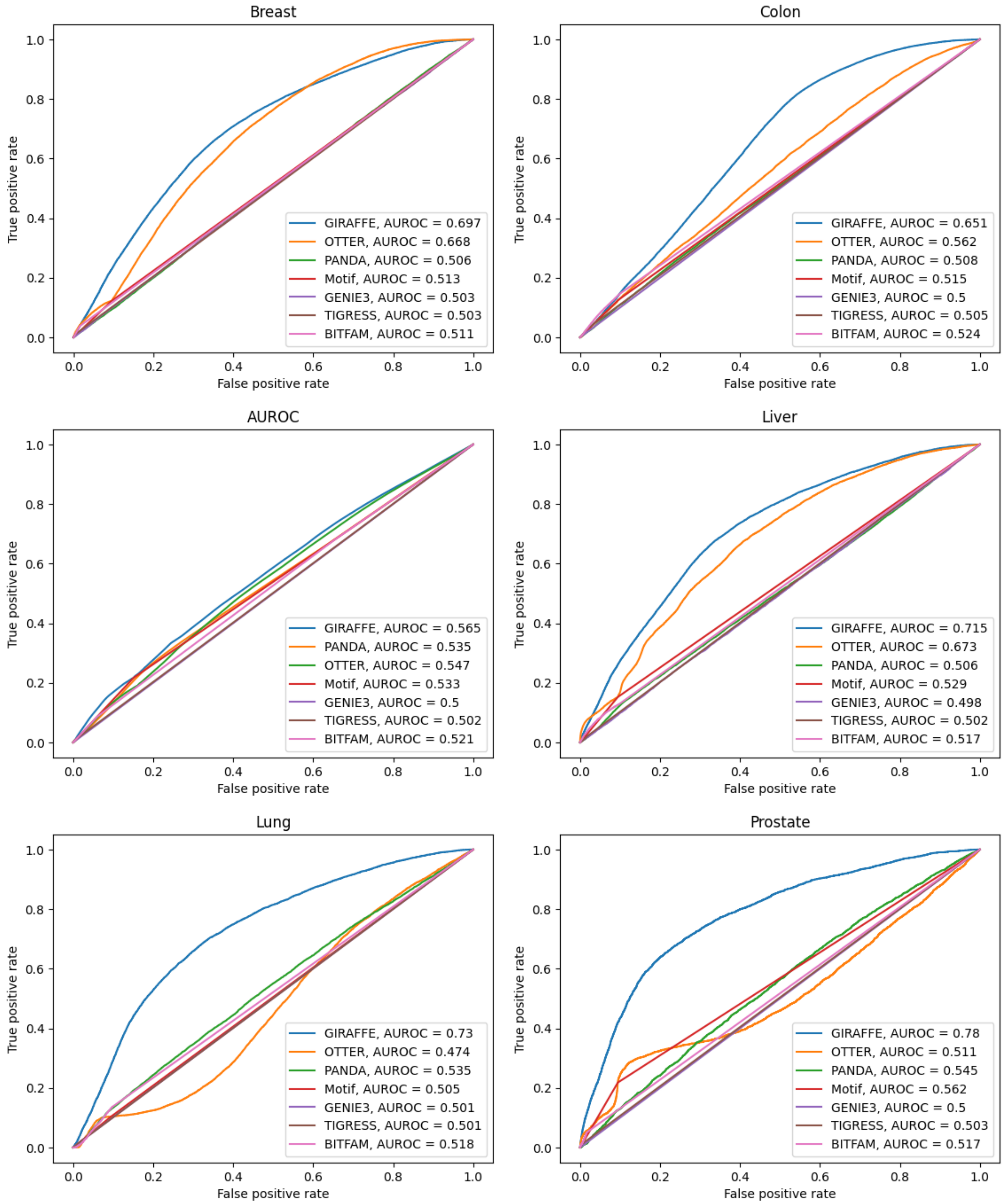


Figure S1: Receiver operating characteristic (ROC) curves for the tissue-specific ChIP-seq validation analyses summarized in Table 2 of the main text. Each panel corresponds to one tissue.

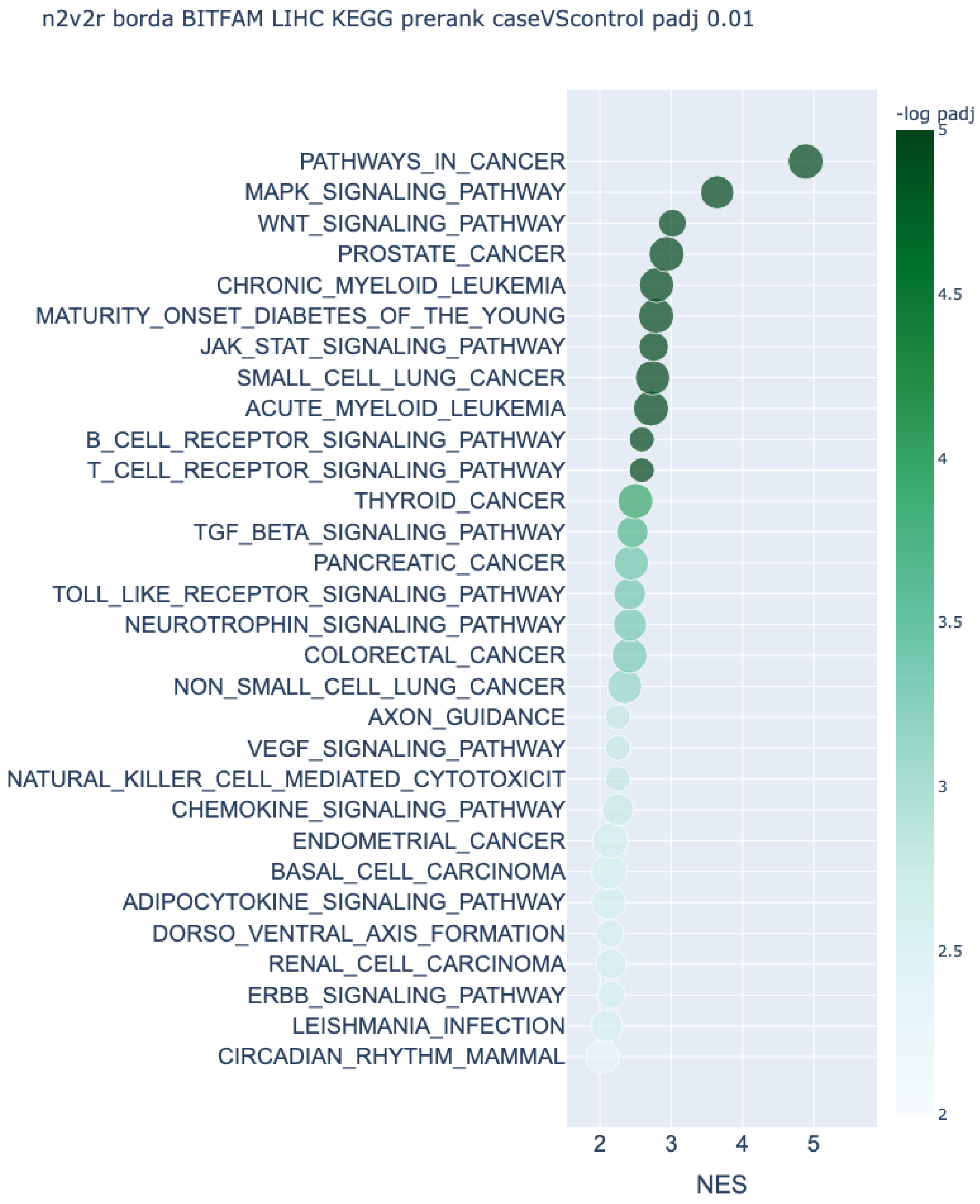


Figure S2: **KEGG pathways identified from differential Bitfam gene regulatory networks between healthy liver and LIHC.** Pathway enrichment was performed using GSEA on node2vec2rank embeddings (Kolmogorov–Smirnov test, Benjamini–Hochberg FDR-adjusted $p < 0.01$). Compared with Giraffe, the enriched pathways are fewer in number and generally less directly related to liver cancer biology.
